# Large-scale hyperparameter search for predicting human brain responses in the Algonauts challenge

**DOI:** 10.1101/689844

**Authors:** Kamila M. Jozwik, Michael Lee, Tiago Marques, Martin Schrimpf, Pouya Bashivan

## Abstract

Image features computed by specific convolutional artificial neural networks (ANNs) can be used to make state-of-the-art predictions of primate ventral stream responses to visual stimuli.

However, in addition to selecting the specific ANN and layer that is used, the modeler makes other choices in preprocessing the stimulus image and generating brain predictions from ANN features. The effect of these choices on brain predictivity is currently underexplored.

Here, we directly evaluated many of these choices by performing a grid search over network architectures, layers, image preprocessing strategies, feature pooling mechanisms, and the use of dimensionality reduction. Our goal was to identify model configurations that produce responses to visual stimuli that are most similar to the human neural representations, as measured by human fMRI and MEG responses. In total, we evaluated more than 140,338 model configurations. We found that specific configurations of CORnet-S best predicted fMRI responses in early visual cortex, and CORnet-R and SqueezeNet models best predicted fMRI responses in inferior temporal cortex. We found specific configurations of VGG-16 and CORnet-S models that best predicted the MEG responses.

We also observed that downsizing input images to ~50-75% of the input tensor size lead to better performing models compared to no downsizing (the default choice in most brain models for vision). Taken together, we present evidence that brain predictivity is sensitive not only to which ANN architecture and layer is used, but choices in image preprocessing and feature postprocessing, and these choices should be further explored.

## Introduction

In recent years, deep convolutional artificial neural networks (ANNs) have revolutionized computer vision and achieved high performance on the ImageNet object recognition challenge. Many studies in the last several years have demonstrated considerable similarities between ANNs and brain representations of objects. Specifically, ANNs predict representations of object images in the visual cortex, as measured in humans via fMRI (e.g. (1)), and MEG (e.g., (2)). Only a small number of ANN architectures were previously tested in their ability to predict neural representation in the human brain (e.g., (1–3)). Here, we assessed the representational similarity of a wide range of network architectures to measurements from human brains given specific networks, input preprocessing strategies, feature pooling mechanisms, and dimensionality reduction approaches.

## Methods

### fMRI

Subjects (n=15, two separate groups) had their brain activity measured with a 3T fMRI scanner while they viewed two sets of images: 92 colored images of real-world objects segmented from their backgrounds and presented on a gray background (data from (4)) and 118 colored images with backgrounds (data from (2)). Images were presented at the center of fixation (size: 2.9° visual angle, stimulus duration: 500 ms for 92 images; and size: 4° visual angle, stimulus duration: 500 ms for 118 images) as the subjects were performing a fixation task. Regions of interest (early visual cortex (EVC) and inferior temporal (IT) cortex) were defined anatomically in each subject.

### MEG

Two groups of subjects viewed the same two sets of images used for fMRI (N= 15 for each image set) while their brain activity was measured with MEG (data from (4) and (2)). The experimental design was similar to the fMRI study described above. Early (70-90 ms for 92 images, and 100-120 ms for 118 images) and late (140-160 ms, and 165-185 ms) time windows were used in the analyses. Peak latency was 82 ms (early) and 150 ms (late) for 92 images and 108 ms (early) and 176 ms (late) for 118 images. For each subject and time window (early and late), we averaged the representational dissimilarity matrices (explained below) computed for each 10ms time window.

### Deep Artificial Neural Networks

We considered 33 ANN architectures from recent neural networks literature (full list of models is available on www.brain-score.org (5). All ANNs were trained to classify 1.2 million images into 1000 categories using the ImageNet dataset (6). We extracted activations from a range of ANNs for the two image sets (described above).

### Grid search

We performed grid search over network architectures and input and output parameterizations to find the models that have the highest correlation with brain representations (Figure 1). Our goal was to assess to what degree these parameters could improve our ability to predict the human brain responses to natural visual stimuli. Specifically, we tested the following parameters: input field of view (to account for the uncertainty over visual extent of input tensor), performing Gaussian blur or adding jitter to images (to account for adversarial regions), averaging activations across filter types (depth average, to account for smoothing over space in fMRI and MEG), averaging activations across the filter map (spatial average, to account for smoothing over space in fMRI and MEG), and applying PCA on ANN activations (to prevent overfitting). We extracted activations for all the models, layers, and parameter combinations yielding more than 140,338 candidates.

**Fig. 1.**
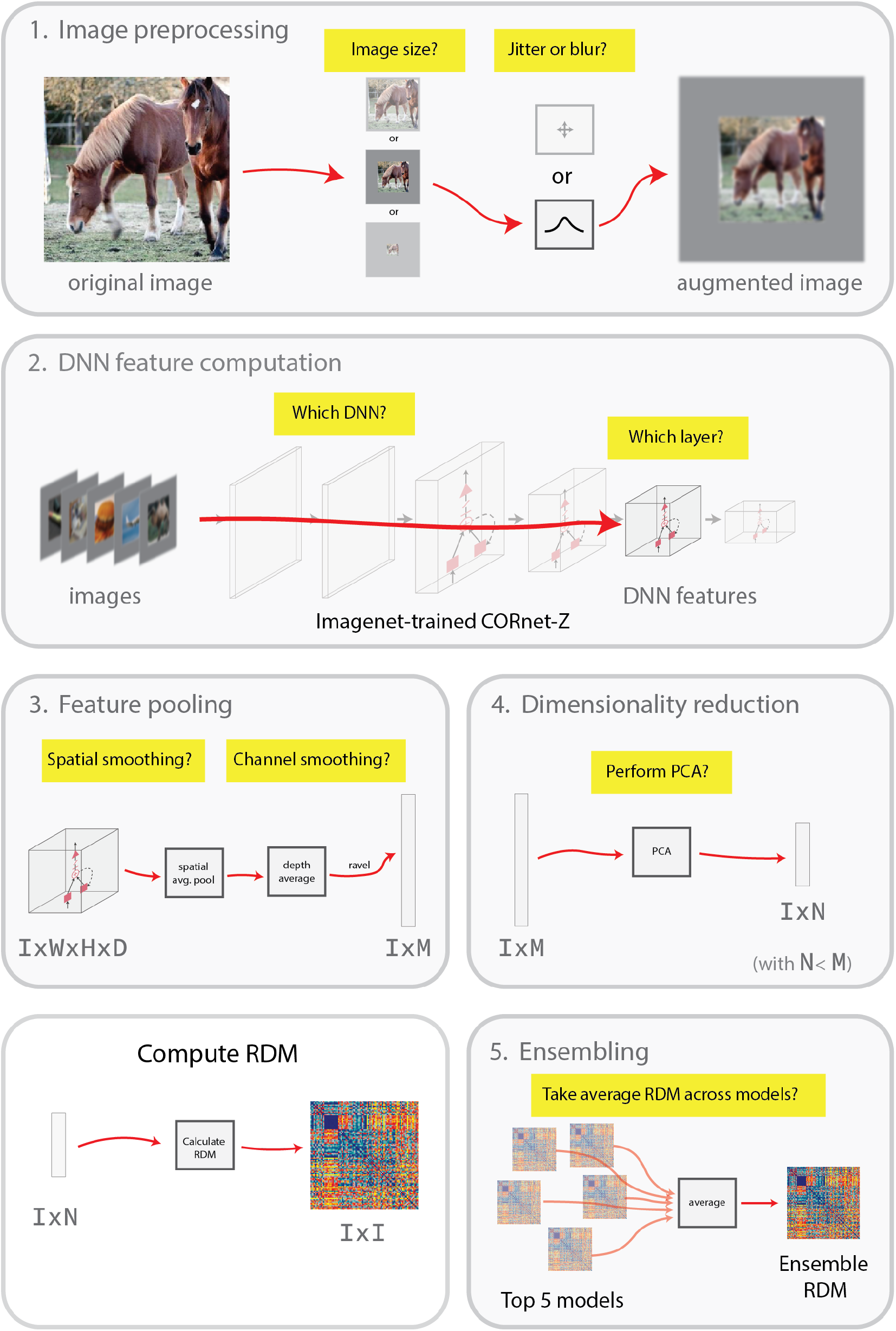
Grid search over parameters. Parameters for image preprocessing included varying the model input field of view and adding or not blur or jitter to the images. We extracted activations for two sets of images (92 and 118 images) from a range of networks and layers after each computational block in a network. We either left the activations as they were or added spatial and/or depth averaging. We either applied, or did not apply (PCA) on the activations. We computed RDMs for activations from each grid condition. Finally, we correlated RDMs with fMRI RDMs (EVC and IT) or MEG RDMs (early and late timepoints) and obtained the similarity scores. We noise corrected the scores for each dataset (92 or 118) images and averaged the scores across the two datasets to obtain a mean score. We combined five RDMs with the best scores into one averaged RDM.

**Table 1.** Best model information. We identified the five best models with the highest average score across the two datasets (92 and 118 images) for each data type (fMRI EVC, fMRI IT, early MEG, and late MEG). The best model and layer names are provided in the table together with their parameter specifications. Parameters of the images included image scale and the presence or absence of jitter and blur. Parameters of the ANN activations included spatial pooling, depth averaging, and applying PCA.

|  | Image Scale | Image Noise | DNN | Layer | Spatial Pooling | Depth Average | PCA | Score (R) |
| --- | --- | --- | --- | --- | --- | --- | --- | --- |
| fMRI EVC | 50% | blur | CORnet-S | V2.output-t0 | to 16x16 | yes | no | **0.3734** |
|  | 50% | blur | CORnet-S | V2.output-t0 | none | yes | no | 0.3722 |
|  | 50% | none | CORnet-S | V2.output-t0 | to 16x16 | yes | no | 0.3711 |
|  | 50% | jitter | CORnet-S | V2.output-t0 | to 16x16 | yes | no | 0.3710 |
|  | 50% | jitter | CORnet-S | V2.output-t0 | none | yes | no | 0.3700 |
| fMRI IT | 50% | blur | SqueezeNet1.1 | features.11 | to 4x4 | no | no | 0.1257 |
|  | 50% | jitter | SqueezeNet1.1 | features.11 | to 4x4 | no | no | 0.1246 |
|  | 75% | blur | CORnet-R | IT.output-t3 | to 4x4 | no | no | 0.1227 |
|  | 75% | blur | CORnet-R | IT.output-t3 | to 8x8 | no | no | 0.1227 |
|  | 75% | blur | CORnet-R | IT.output-t3 | to 16x16 | no | no | 0.1277 |
| MEG (early) | 50% | jitter | VGG-16 | block3_pool | to 16x16 | no | no | 0.4796 |
|  | 50% | none | VGG-16 | block3_pool | to 16x16 | no | no | 0.4779 |
|  | 50% | blur | VGG-16 | block3_pool | to 16x16 | no | no | 0.4752 |
|  | 50% | blur | CORnet-S | V2.output-t0 | to 16x16 | yes | no | 0.4730 |
|  | 50% | jitter | CORnet-S | V2.output-t0 | to 16x16 | yes | no | 0.4722 |
| MEG (late) | 75% | blur | CORnet-S | V2.output-t0 | to 16x16 | yes | no | 0.4488 |
|  | 50% | none | CORnet-S | V2.output-t0 | to 16x16 | yes | no | 0.4378 |
|  | 75% | blur | CORnet-S | V2.output-t0 | none | yes | no | 0.4366 |
|  | 50% | jitter | MobileNet v1 | Conv2d 3 | none | no | no | 0.4363 |
|  | 50% | jitter | CORnet-S | V2.output-t0 | to 16x16 | yes | no | 0.4359 |

### Representational Similarity Analysis

We computed response patterns (based on fMRI, MEG, and ANNs) for each image. We then computed response-pattern dissimilarities between each pair of images and placed these in a representational dissimilarity matrix (RDM). An RDM captures which distinctions among stimuli are emphasized and which are de-emphasized by a particular model or brain region. We estimated model performance by correlating model and data RDMs using Spearman correlation.

## Results

### Track 1: fMRI responses

We assessed the ability of ANNs to predict EVC and IT fMRI responses for two datasets (92 and 118 images). The model and data similarity scores for over 140,338 parameter combinations were spread and we selected the models with the highest similarity score with brain data (Figure 2). Our goal was to find the best model that could predict the brain responses regardless of the stimulus set. Therefore we selected models with high similarity scores across the two datasets. Among the first five models that have high EVC predictivity and are common for the two datasets is CORnet-S (Figure 3). CORnets (7) have an architecture that approximates the number and size of visual areas in the macaque brain. CORnet-S is a recurrent ANN with skip connections. For IT, models with good predictivity are SqueezeNet v1.1 and CORnet-R. SqueezeNet is an ANN with AlexNet-level accuracy but with 50x fewer parameters.

**Fig. 2.**
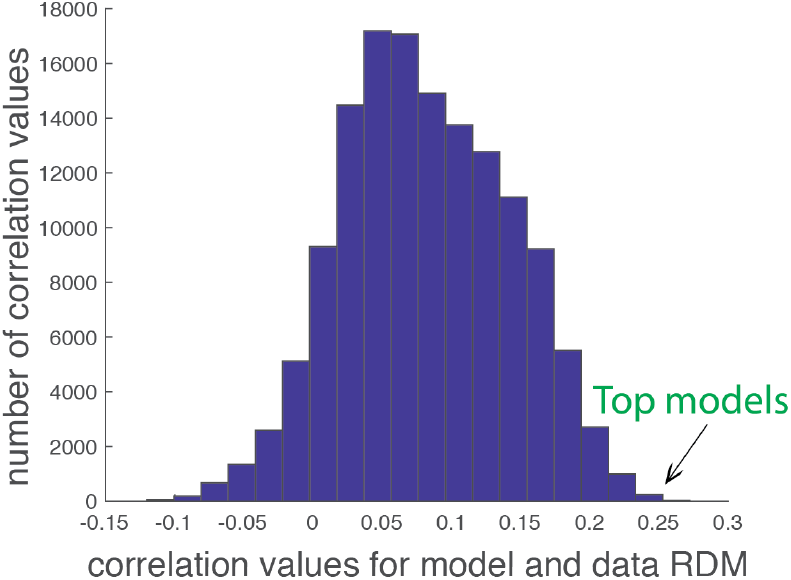
Distribution of correlation values for model and data RDMs for grid search conditions for predicting IT (92 images).

### Track 2: MEG responses

We correlated each model with early and late time windows in MEG responses. Averaged early MEG responses are best predicted by VGG-16 and CORnet-S. Averaged late MEG responses are best predicted by CORnet-S and MobileNet v1 1.0 160. MobileNets are light-weight ANNs whose architecture uses depth-wise separable convolutions.

### Trends in the most predictive models

We observed some qualitative trends in the top performing models, and report them informally here:

- The best input field of view for most of the top models is 0.5, meaning the images occupy half of the models’ inputs.
- Blur or jitter added to the images seem to be beneficial.
- All models that best predict EVC have depth averaging.
- PCA does not seem to be beneficial.

We combined the five best model RDMs and averaged them, which resulted in one combined model per dataset. We used these models to make a single submission to the Algonauts challenge (8), a competition aimed at finding models that best predict human fMRI and MEG brain representation.

## Discussion

In summary, our results suggest that there is a certain class of models that is consistently among the best models that predict brain responses in fMRI and MEG - CORnets. We also found that priors on image size and presence of blur or jitter in images seem to have a positive effect on the predictivity of brain representations. A more detailed analysis is needed to determine to what extent each of the tested parameters beyond network architectures (input preprocessing strategies, feature pooling mechanisms, and dimensionality reduction approaches) contributes to predicting brain representations. It is possible that some of the parameters may be more relevant to certain brain areas but not others (e.g., depth averaging seemed to be beneficial for EVC but not IT). Our approach could be useful to define a set of parameters to be used to predict representations in a given brain area or modality, establishing a new ANN processing pipeline in the field.

## ACKNOWLEDGEMENTS

This work was funded by the Sir Henry Wellcome Postdoctoral Fellowship (206521/Z/17/Z) to KMJ, the Simons Foundation grants (SCGB [325500, 542965]) to JJD, the Semiconductor Research Corporation (SRC) to MS and MJL, and the Center for Brains, Minds and Machines (NSF STC award CCF-1231216).

